# A meta-analysis of Microarray Data is Effective for Identifying Gravity-Sensitive Genes

**DOI:** 10.1101/2021.03.31.437982

**Authors:** Yin Liang, Mengxue Wang, Yun Liu, Chen Wang, Ken Takahashi, Keiji Naruse

## Abstract

Gravity affects the function and maintenance of organs, such as bones, muscles, and the heart. Several studies have used DNA microarrays to identify genes with altered expressions in response to gravity. However, it is technically challenging to combine the results from various microarray datasets because of their different data structures. We hypothesized it is possible to identify common changes in gene expression from the DNA microarray datasets obtained under various conditions and methods. In this study, we grouped homologous genes to perform a meta-analysis of multiple vascular endothelial cell and skeletal muscle datasets. According to the t-distributed stochastic neighbor embedding (t-SNE) analysis, the changes in the gene expression pattern in vascular endothelial cells formed specific clusters. We also identified candidate genes in endothelial cells that responded to gravity. Further, we exposed human umbilical vein endothelial cells to simulated microgravity using a clinostat and measured the expression levels of the candidate genes. Gene expression analysis using qRT-PCR revealed that the expression level of the prostaglandin transporter gene *SLCO2A1* decreased in response to microgravity, consistent with the meta-analysis of microarray datasets. Furthermore, the direction of gravity affected the expression level of *SLCO2A1*, buttressing the finding that its expression was affected by gravity. These results suggest that a meta-analysis of DNA microarray datasets may help identify new target genes previously overlooked in individual microarray analyses.

## 1 Introduction

Long-term space missions, such as those to Mars, have become more feasible. During space missions, astronauts experience microgravity, which is the condition of being weightless. During spaceflight, the astronauts’ cardiovascular function undergoes tremendous changes, such as blood redistribution, increased compliance with lower extremity veins, heart contraction, and orthostatic intolerance [1, 2]. Thus, it is crucial to study the impact of spaceflight on various organs, such as the cardiovascular system. A recent study on twin astronauts demonstrated that a one-year spaceflight induced various changes, such as carotid artery distension and increased intima-media thickness [3]. This study suggests that retinal edema formation in space may be promoted by angiogenesis. Therefore, it is vital to elucidate the effect of microgravity on vascular endothelial cells for the coming space age.

Microgravity affects the morphology and physiology of the cardiovascular system. While the microgravity-induced changes are dramatic, they are thought to be mostly secondary to headward fluid shifts rather than direct effects of microgravity on the cardiovascular system [4]. Another factor that microgravity exerts on the orthostatic intolerance is a decrease in constrictor responses of mesenteric artery and vein [5].

On the other hand, the endothelium is one of the most sensitive tissues to changes in gravity [6, 7]. Microgravity directly acts on vascular endothelial cells and affects processes, such as angiogenesis [8]. For example, microgravity causes the down-regulation of actin [9] and cytoskeletal disorganization [10] and decreases the metabolism of the endothelial cells [11].

Several models, such as human subjects in bed rest, the suspended hind limbs of a rodent, clinorotated cultured cells, and humans, animals, and cultured cells in spaceflight [12-14], are used to evaluate the effect of microgravity on organs. On the other hand, DNA microarray is often performed using RNA extracted from various tissues to analyze the gene expressions affected by microgravity.

However, the studies examining the effect of microgravity on gene expression tend to yield inconsistent results. Although many investigations generate gene expression data, only a few have cross-sectionally examined data, especially in space biology and medicine. Two factors make such comparisons of gene expression among multiple datasets difficult. First, the studies used different microarrays to analyze different gene sets. Second, the nomenclatures of the same genes differ in the studies.

We hypothesized that it was possible to identify common, essential microgravity-induced changes in gene expression from the results obtained using varying materials and methods. This study aimed to uncover the overlooked effects of microgravity on gene expression from accumulated valuable database resources by using different statistical methods. In this study, we applied a method developed by Kristiansson et al. [15] to categorize genes into homologous groups and perform a meta-analysis to resolve the problem of genes having different names in various DNA arrays. Since microgravity causes not only cardiovascular deconditioning but also muscle atrophy, we analyzed the changes in gene expression in endothelial cells and skeletal muscle obtained from microarray datasets in multiple simulated or in-flight microgravity experiments.

It would also be fascinating and important to identify the similarity in the impact of microgravity on the gene expression in different species. However, due to the variety of materials and methods used, it is difficult to perform a consistent analysis of the genes that respond to microgravity. To our knowledge, the similarity between the results from gene expression analysis obtained with different materials and methods has never been shown systematically in space biology. Here, we demonstrated such similarity by applying the t-distributed stochastic neighbor embedding (t-SNE) analysis [16], a method extensively used for visualizing high-dimensional data from flow cytometry to DNA array datasets.

Furthermore, after the gravity-sensitive genes were identified from the meta-analysis, their change in expression in human umbilical vein endothelial cells (HUVECs) was induced by simulated microgravity (SMG) using a clinostat and tested with qRT-PCR.

We found that the levels of *LOX, SLCO2A1*, and *TXNIP* expression were significantly altered in response to SMG. Also, we found that the direction of gravity affected the expression of *SLCO2A1*. Instead of being specific to the condition of microgravity, the expression of *SLCO2A1* may be altered by gravity from a direction different from the arrangement of its cytoskeleton. These results suggest that the meta-analysis of accumulated microarray data is useful for identifying candidate genes with altered expression in response to gravity changes. This strategy could potentially contribute to the advancement of space biology and medicine.

## 2 Materials and methods

### 2.1 Microarray analysis

The gene expression data for endothelial cells were obtained from two studies, Gene Express Omnibus (GSE43582) [7] and Ma et al. [17] (Table 1), which investigated the effects of microgravity. Ma et al. made two comparisons. It compared the 1 G and SMG adherent groups, and between the 1 G and SMG three-dimensional (3D) aggregate groups on day 7. Then, we performed a meta-analysis to test the null hypothesis that a gene of interest was not differentially expressed among all genes in the databases using a method described by Kristiansson et al. [15]. The cross-experimental p-values were corrected for multiple testing using the false discovery rate (FDR) determined via the Benjamini-Hochberg procedure. For the muscle samples, gene expression data from 12 experiments investigating the effects of microgravity were obtained from Gene Express Omnibus (Table 2). We performed the same meta-analysis and p-value correction for endothelial cells.

### 2.2 t-SNE analysis

A t-SNE analysis was performed using R’s tsne package (https://cran.r-project.org/web/packages/tsne/) to group the multiple DNA array datasets. The perplexity parameter in the t-SNE analysis indicates the effective number of neighbors, and the result of the t-SNE plot may differ depending on its setting. The grouping by t-SNE analysis is shown to the same regardless of the value of perplexity by performing the t-SNE analysis with varying perplexity parameter 1, 2, or 3. In this study, the t-SNE analysis was performed at least 10 times at each perplexity parameter to confirm that the results showed a constant pattern. Furthermore, cluster analysis was performed using cluster 3.0 to group multiple DNA array datasets by a method different from the above t-SNE analysis [18].

### 2.3 HUVEC culture

The HUVECs (Lonza, Basel, Switzerland) were cultured in PromoCell Endothelial Cell Growth Medium 2 containing 0.02 ml/ml fetal calf serum. In a T-25 culture flask (Cat. #: 353107, Corning, NY, USA), 1 × 10^5^ cells were cultured at 37°C at 5% CO_2_ for the duration of the experiment.

### 2.4 Exposure to SMG

The HUVECs were seeded in the flasks and cultured for 2 days at 37°C at 5% CO_2_ to ensure the cells attach to the bottom of the flasks. Then, the flasks were split into two groups, SMG and 1 G control. The SMG flasks were clinorotated for 7 days using a clinostat (Zeromo; Kitagawa Iron Works Co. Ltd., Hiroshima, Japan) to achieve 10^−3^ G in the incubator (Fig. 4A). During clinorotation, air bubbles were removed from the flasks to prevent shear stress. The 1 G control flasks were placed next to the clinostat and kept static for 7 days.

### 2.5 Cell culture under different gravitational configurations

The HUVECs were seeded in the flasks simultaneously and cultured for 2 days to confirm the cells’ attachment. Then, the flasks were divided into the “up,” “side,” and “down” configures, where the cells were on a horizontal surface, on a vertical surface, and hung on a horizontal surface, respectively (Fig. 7A). The “up” configuration was the control condition. This experiment was performed independently from the clinostat experiment in Section 2.4.

### 2.6 Phase-contrast images of HUVECs

For the SMG experiments, the cells’ condition was observed by acquiring daily phase-contrast images of the same areas in the culture flask using a TE2000 inverted microscope. For the experiment involving different gravitational configurations, the same areas in the culture flask were imaged on the day before exposure to the gravitational conditions and after 7 days of exposure to the conditions to observe the morphology and growth of the cells.

### 2.7 qRT-PCR

Total RNA was harvested from cells using a High Pure RNA Isolation Kit (Roche Diagnostics Indianapolis, IN, USA), according to the manufacturer’s protocols. The RNA concentration was measured using a spectrometer (Nanodrop ND-1000, Thermo Fisher Scientific, USA). RNA samples with an OD 260/280 ratio of more than 1.80 were used for subsequent procedures. The samples were stored at −80°C.

For qRT-PCR, RNA samples were first heated at 70°C for 5 min to destroy the secondary structures. Then, the 1 ng of RNA was reverse-transcribed into cDNA using a Verso cDNA Synthesis Kit (Thermo Scientific, Lithuania, USA). The cDNA was used for qRT-PCR to quantify the expression of 20 genes (Table 3) using forward and reverse primers from previous publications or designed using the Primer3 (Version 0.4.0, http://bioinfo.ut.ee/primer3-0.4.0/) software. SYBR Green reagents (Life Technologies) were used for the relative quantification of RNA levels according to the manufacturer’s instructions. Lastly, 18S rRNA was used as an internal control.

### 2.8 Statistical analysis

Statistical analysis was performed using Prism (version 6.07, GraphPad Software, La Jolla, CA, USA). The data were expressed as mean ± SD. The significance of the differences between the experimental groups was determined using the Student’s t-test or Kruskal-Wallis test with Dunn’s Multiple Comparison. A p-value of 0.05 or less was considered statistically significant.

## 3 Results

### 3.1 Statistical meta-analysis of gene expression in endothelial cells in response to microgravity conditions

We analyzed the gene expression data from microgravity exposure experiments performed in spaceflight or with clinorotation in the laboratory (Table 1) to identify genes in endothelial cells with altered expression in response to microgravity. A total of 20,358 genes were retrieved from the spaceflight experiment, and 903 genes were retrieved from the clinorotation experiment. At first, GEO2R [19] was used to identify the genes differentially expressed in the ground control and flight samples. The p-values obtained from this analysis were used with the p-values from the comparison between the 1 G and clinorotated adherent groups provided by Ma et al. [X] for the subsequent meta-analysis. Applying the method described by Kristiansson et al. [X], 108 genes with a p-value of 0.001 or less were identified, of which 46 genes had a FDR of 0.05 or less (Fig. 1A). We randomly chose 10 genes from the 46 genes for subsequent gene expression assays using qPCR. Of the 10 genes, *CCN5*, encoding WISP2 (WNT1 Inducible Signaling Pathway Protein 2), had the lowest p-value of 2.13 × 10^−8^, also the point with a high −log_10_FDR value in the volcano plot (Figs. 1B and 1C). *LOX* (encoding lysyl oxidase) and *MT1X* (encoding metallothionein 1X) showed higher fold change values (logFC) at 4.11 and 1.71, respectively, in clinorotated EA.hy926 cells, indicating that microgravity facilitated the changes in their expression (Fig. 1C). However, the logFC values of *LOX* and *MT1X* were −0.35 and −0.45, respectively, in the space-flown HUVECs, indicating the decreased expression of these genes. The logFC value of *SLCO2A1* (encoding solute carrier organic anion transporter family member 2A1) was −2.10 in the clinorotated EA.hy926 cells and −0.23 in the space-flown HUVECs, with an FDR of 5.18 × 10^−2^.

**Figure 1.**
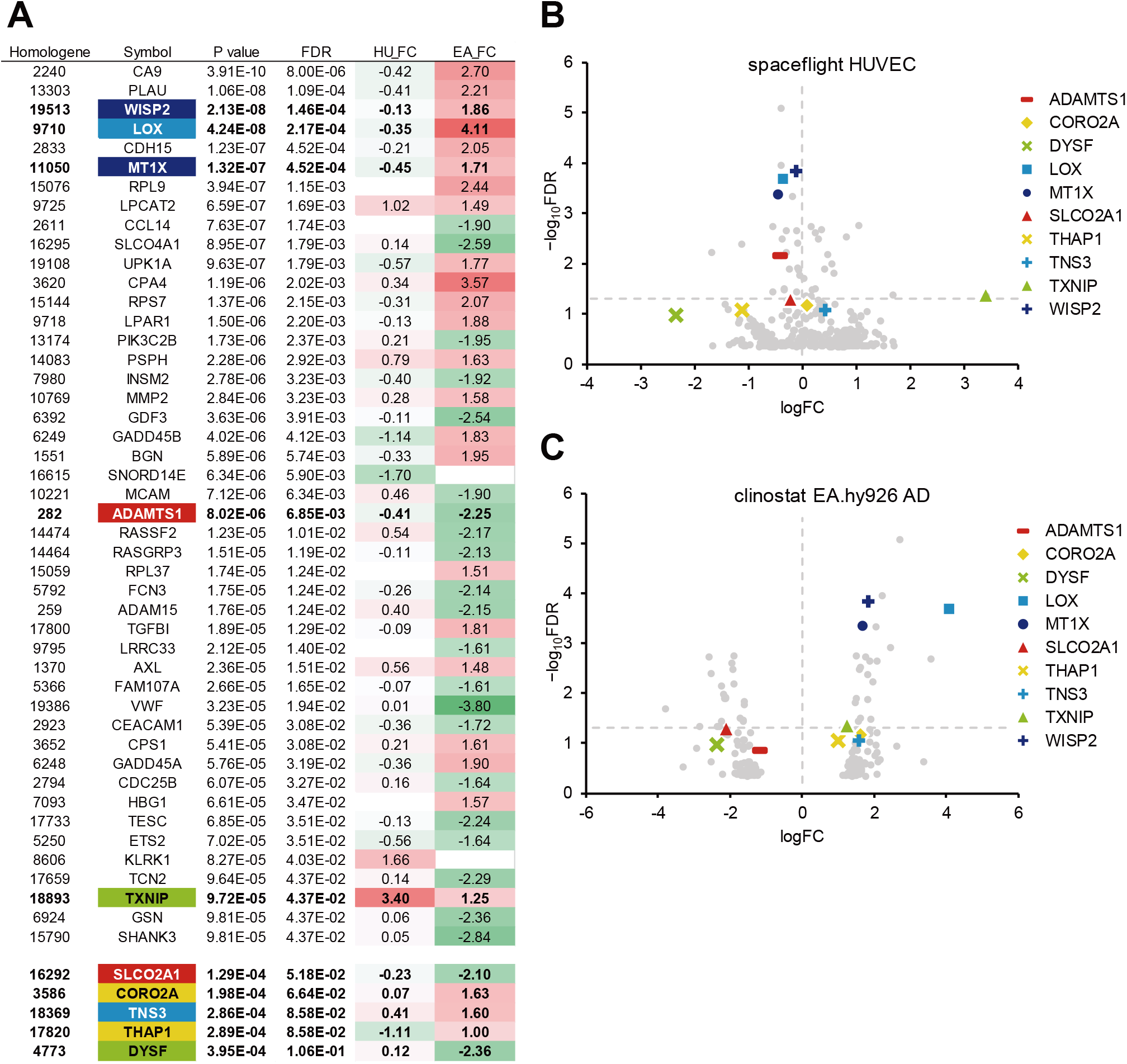
A meta-analysis of gene expression in endothelial cells in response to microgravity conditions. (A) Top 46 genes with the smallest p-values (FDR < 0.05), which were identified from the DNA array analyses of human umbilical vein endothelial cells (HUVEC) and EA.326hy cells under spaceflight and simulated microgravity conditions, respectively. Additional genes with relatively low p-values were listed below the top 46 genes. HU_FC: Log fold change value in HUVEC. EA_FC: Log fold change value in EA.hy926 cell. (B) The Volcano plot of the space flight HUVECs. (**C**) The Volcano plot of the clinorotated EA.hy926 cells. Horizontal dotted line: a line at which FDR is equal to 0.05. Vertical dotted line: a line at which logFC is equal to zero.

Then, we performed another meta-analysis using the same p-values from the space-flown HUVECs and those from another microgravity condition, the 3D aggregates in clinorotated EA.hy926 cells. As a result, 79 genes with a p-value of 0.001 or less were identified, of which 64 genes had a FDR of 0.05 or less (Fig. 2A). Again, *LOX* showed a significant p-value of 2.95 × 10^−7^ and a high logFC value of 3.44 in the clinorotated EA.hy926 cells, with the logFC of−0.35, in opposite direction in the space-flown HUVECs (Figs. 2A-C). In contrast, *SLCO2A1* showed a significant logFC value of −2.66 and an FDR of 2.60 × 10^−3^, with a logFC of −0.23, in the same direction in the space-flown HUVECs.

**Figure 2.**
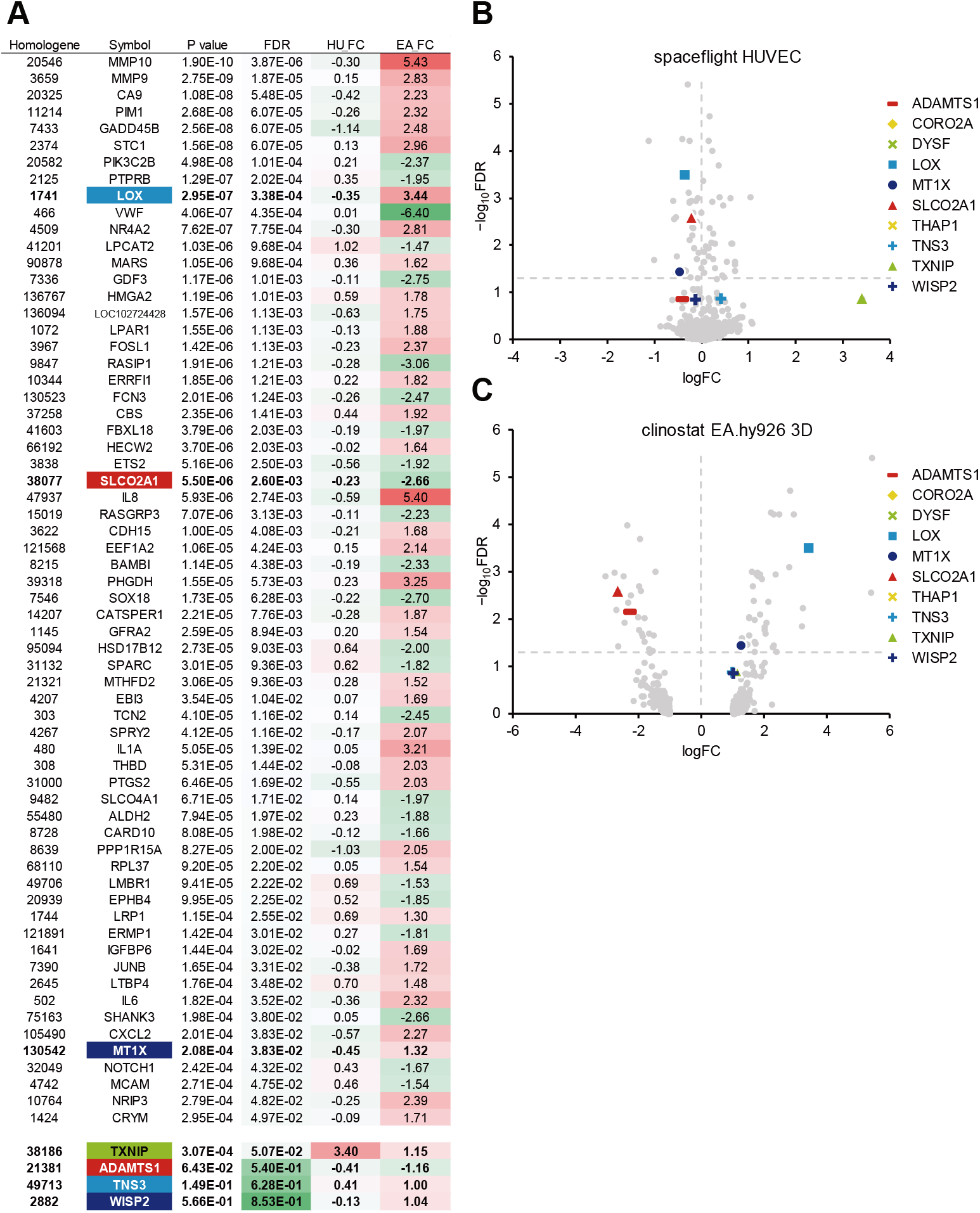
A meta-analysis of gene expression in endothelial cells in response to other microgravity conditions. (A) Top 64 genes with the smallest p-values (FDR < 0.05), which were identified from the DNA array analyses of human umbilical vein endothelial cells (HUVEC) and EA.326hy cells under spaceflight and simulated microgravity conditions, respectively. Additional genes with relatively low p-values were listed below the top 46 genes. HU_FC: Log fold change value in HUVEC, EA_FC: Log fold change value in EA.hy926 cell. (B) The Volcano plot of the space flight HUVECs. (C) The Volcano plot of the clinorotated EA.hy926 cells. Horizontal dotted line: a line at which FDR is equal to 0.05. Vertical dotted line: a line at which logFC is equal to zero.

### 3.2 Statistical meta-analysis of gene expression in skeletal muscle in response to microgravity conditions

Next, we analyzed the gene expression data from the experiments using multiple microgravity loads on skeletal muscle (Table 2). The p-values were obtained using GEO2R from each dataset. For example, the differential expression of genes across pre and post-bed rest experiments in the soleus muscle was analyzed using the GSE14798 dataset. A meta-analysis of 12 datasets identified 6,328 genes with a FDR of 0.05 or less, of which 10 genes exhibited changes in expression in the same direction (both increasing or decreasing) in most of the datasets with valid fold change (FC) values (Fig. 3). These genes were selected for the subsequent gene expression assays using qPCR.

**Figure 3.**
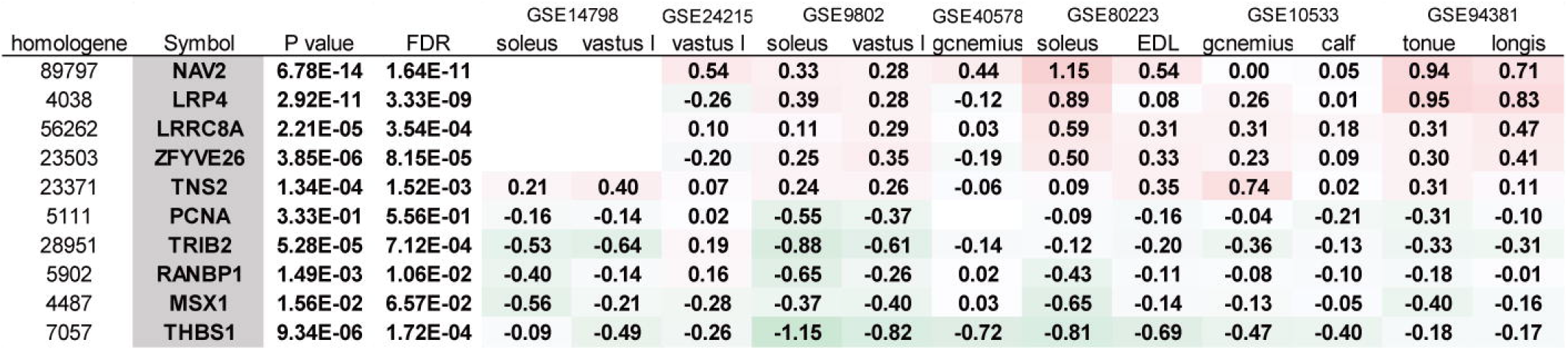
A meta-analysis of gene expression in muscle samples in response to microgravity conditions. The 10 genes exhibiting changes in expression in the same direction (all increasing or decreasing) in most of the datasets were listed. vastus l: vastus lateralis. gcnemius: gastrocnemius. EDL: extensor digitorum longus. longis: longissimus dorsi.

Then, we performed the t-SNE analysis to gain insight into the similarity between the DNA array datasets. We analyzed 3,290 genes with valid FC values in all the datasets listed in Table 2. The individual datasets were marked A–L. The cost function in the t-SNE analysis reflected the difference between the distance of each gene in multidimensions, 3,290 dimensions in this case, and that in 2D. After performing the t-SNE analysis, the cost value converged between 1,000 and 5,000 steps (Fig. 4A). In step 10,000, the cost value converged to 0.480, 0.293, and 0.259 when the perplexity parameter was 1, 2, and 3, respectively. When the perplexity was 1, the array datasets were divided into two groups, i.e., A/B/C/D/E/I/J and F/G/H/K/L (Fig. 4B). A similar tendency was observed when the perplexity was 2 and 3 (Figs. 4C and 4D). Furthermore, a similar pattern was observed in the results of the cluster analysis (Fig. 4I).

**Figure 4.**
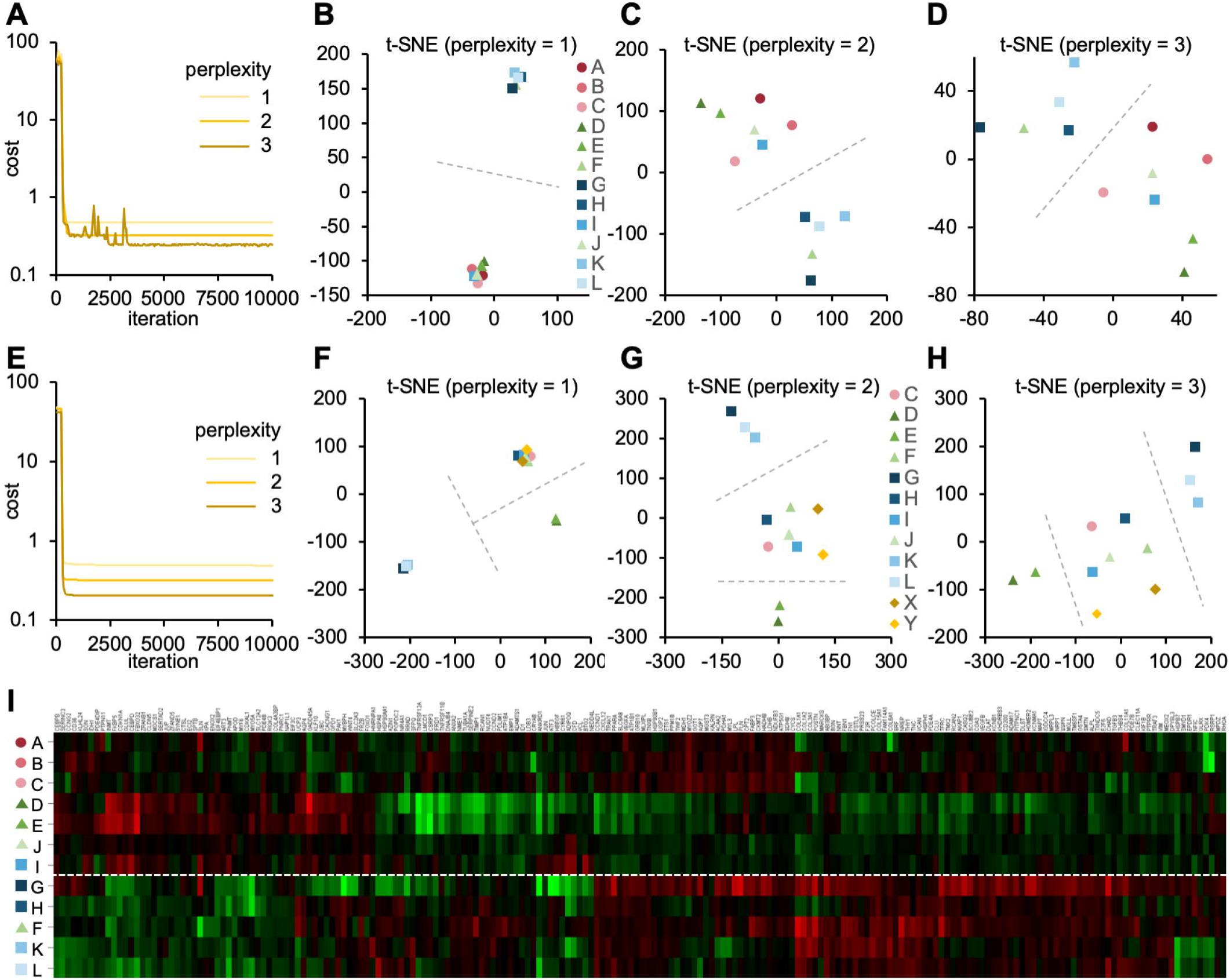
The contribution of experimental conditions to the gene expression pattern. (A–D) The t-SNE analysis of the DNA array datasets obtained from skeletal muscle (Table 2). (A) The convergence of cost values, reflecting the difference between the distance of each gene in multidimensions and that in 2D, over time at the perplexity of 1, 2, and 3. The t-SNE plots at perplexity = 1 (B), 2 (C), and 3 (D) are shown. (E–H) The t-SNE analysis of DNA array datasets obtained from endothelial cells (Table 1) and skeletal muscle (Table 2). (E) The convergence of cost values over time at the perplexity of 1, 2, and 3. The t-SNE plots at perplexity = 1 (F), 2 (G), and 3 (H) are shown. Dashed lines are drawn to guide the grouping of datasets. (I) The cluster analysis of DNA array datasets obtained from skeletal muscle (Table 2).

It would also be interesting to see how the microgravity-induced changes in the gene expression pattern in endothelial cells differ from those in muscle tissue. We investigated this by attempting to perform the t-SNE analysis by combining the 2 array datasets obtained from the endothelial cells (“X” and “Y” in Table 1) with the 12 array datasets obtained from skeletal muscle (“A–L” in Table 2). However, only 52 genes with valid FC values were identified in the combined datasets with 14 arrays. The removal of datasets A and B from the combined datasets resulted in the identification of 177 genes with valid FC values. We conducted the t-SNE analysis using these 177 genes. As a result, the cost value converged within 700 steps (Fig. 4E). In step 10,000, the cost value converged to 0.488, 0.321, and 0.206 when the perplexity was 1, 2, and 3, respectively. When the perplexity was 1, the array datasets were divided into 3 groups: C/F/H/I/J/X/Y, D/E, and G/K/L (Fig. 4F). A similar pattern was observed when the perplexity was 2 and 3 (Figs. 4G and 4H); the endothelium-derived datasets X and Y were relatively close to each other and within the same group.

### 3.3 Morphological analysis of HUVECs under SMG using a clinostat

The shape of the cells in the SMG was compared to those in the 1 G control groups after exposure to gravitational conditions. The groups exhibited similar cell shapes. Also, the cell numbers in the groups were similar, with 622.33 ± 264.66 in the SMG group and 563 ± 101.60 in the 1 G control group (p > 0.05) (Fig. 5B-D).

**Figure 5.**
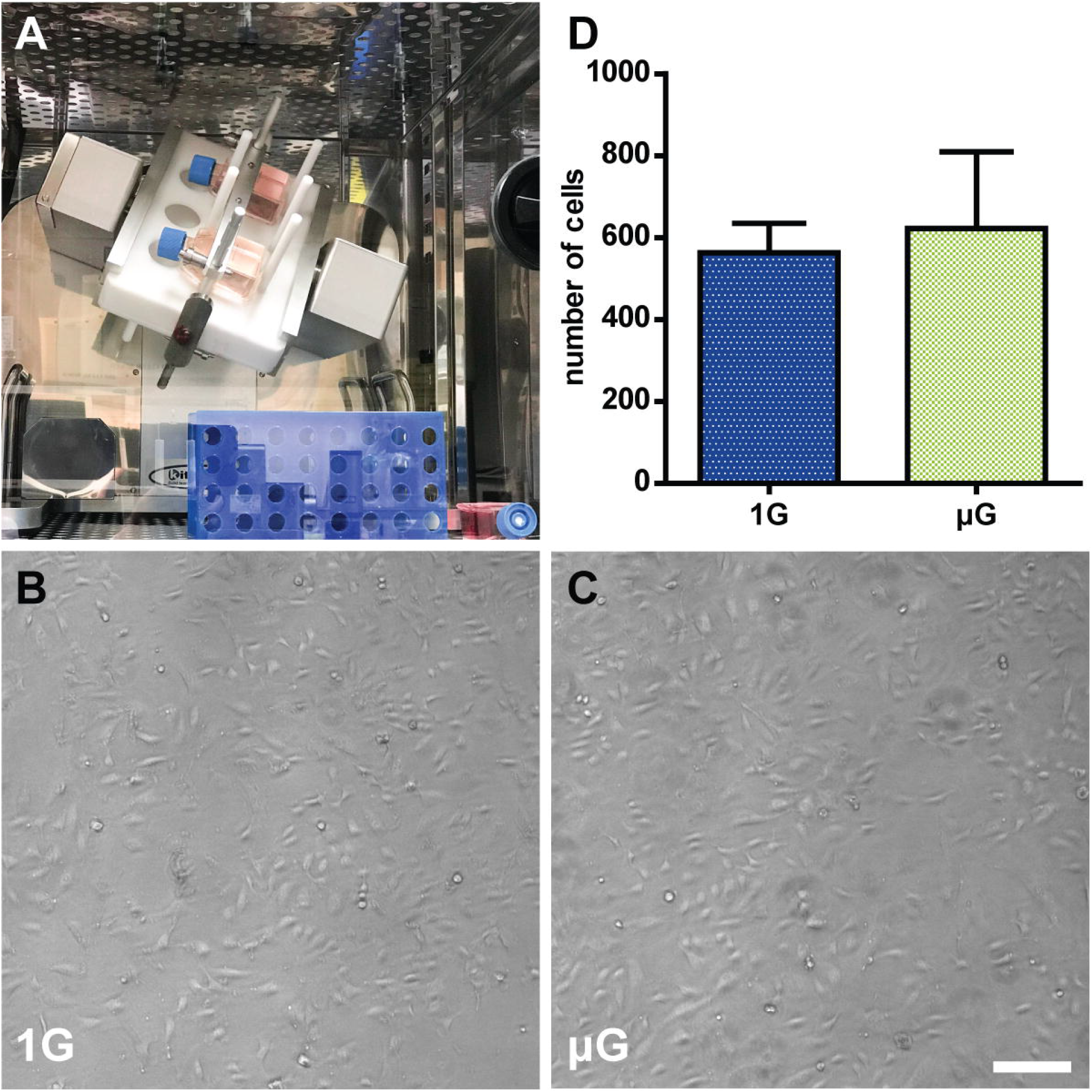
The morphological analysis of human umbilical vein endothelial cells (HUVECs) under simulated microgravity. (A) The clinostat is installed in a CO_2_ incubator. The 1G control flasks were placed next to the clinostat (bottom right). Phase-contrast images of HUVECs after 3 days of exposure to 1 G (B) and simulated microgravity (C). Marker: 200 µm. (D) The number of cells in the simulated microgravity conditions compared to those in the control condition after 7 days of culturing. Data were expressed as the mean ± S.E.M.

### 3.4 Comparison of mRNA expression between control and SMG conditions by qRT-PCR

After exposing the HUVECs to different gravity conditions, we measured the mRNA level of the 10 genes selected from the DNA array meta-analysis of endothelial cells (Fig. 6A). *LOX* expression was higher in the SMG group than in the 1 G group at 1.91 ± 0.27 and 1.20 ± 0.16, respectively (p < 0.05). Conversely, *SLCO2A1* (SMG, 0.43± 0.12; 1G, 0.94 ± 0.05; p < 0.01) and *TXNIP* expression (SMG, 0.43 ± 0.10; 1G, 0.96 ± 0.06; p < 0.01) was lower under the SMG condition than the 1 G condition.

**Figure 6.**
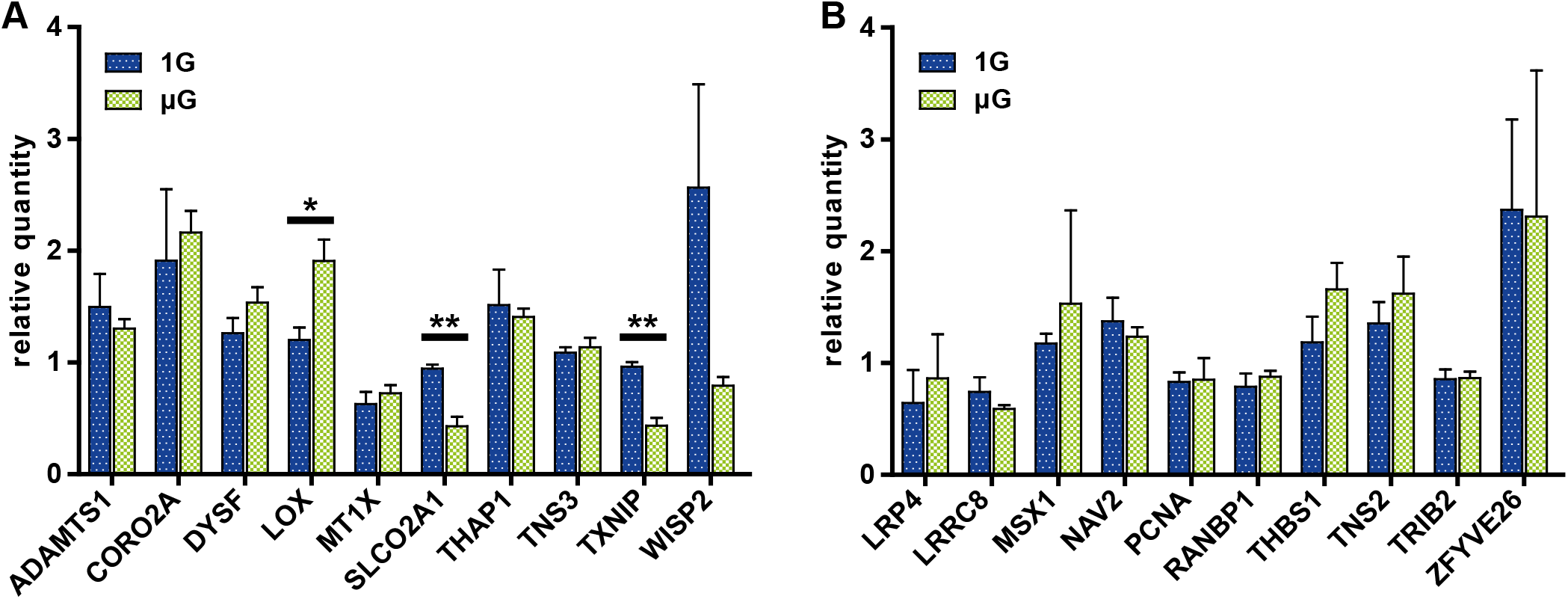
The comparison of gene expression between the control and simulated microgravity conditions in human umbilical vein endothelial cells. (A) The gene sets were obtained from the meta-analyses of datasets from endothelial cells. (B) The gene sets were obtained from the meta-analyses of datasets from muscle samples. The relative quantity of each gene was measured using qRT-PCR and normalized to the level of 18S rRNA. The data were expressed as the mean ± S.E.M. *: p < 0.05, **: p < 0.01.

Next, we measured the expression of the 10 genes selected from the DNA array meta-analysis of muscle tissue (Fig. 6B). In contrast to the result from genes selected from the meta-analysis of endothelial cell DNA arrays, there were no statistically significant differences in gene expression between the SMG and 1 G groups.

### 3.5 Morphological analysis of HUVECs under different gravitational configurations

We tested the effects of the direction of gravity on the morphology and growth of endothelial cells by incubating HUVECs in flasks at the up, side, or down configuration (Fig. 7A). In the down configuration, the cells were less dense than the cells in the up and side configurations (Figs. 7B–D). Moreover, the cells in the down configuration appeared “brushed” and “branched” (Fig. 7D). The number of cells was the smallest in the down configuration at 209.4 ± 112.6 and largest in the side configuration at 638.6 ± 349.4 (Fig. 7E). The difference in cell numbers between the down and side configurations was statistically significant (p < 0.05).

**Figure 7.**
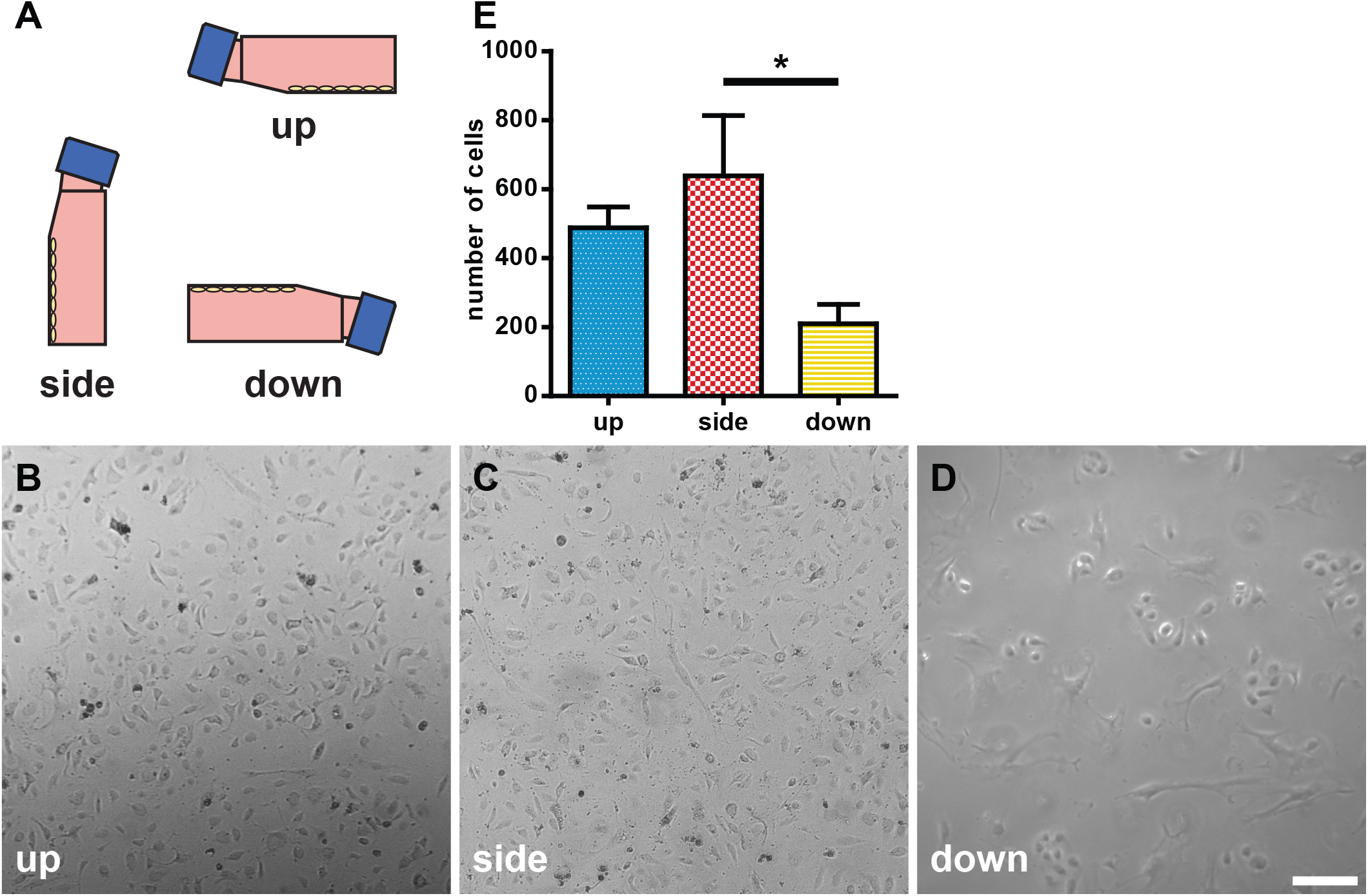
Morphological analysis of human umbilical vein endothelial cells (HUVECs) under different gravitational configurations. (A) The gravitational configurations of cell culture. Left, up configuration; middle, side configuration; and right, down configuration. The phase-contrast images of HUVECs under the up (B), side (C), or down (D) configuration on day 7. Marker: 200 µm. (E) The number of cells in the up, side, and down configurations. The data were expressed as the mean ± S.E.M. *: p < 0.05.

### 3.6 Comparison of mRNA expression among the up, side, and down configurations by qRT-PCR

After exposing HUVECs to different gravity configurations, we measured the expression of the 10 genes selected from the DNA array meta-analysis of endothelial cells (Fig. 8). Kruskal-Wallis test revealed that gravity configuration affected the expression of *SLCO2A1*, with a statistically significant difference (Dunn’s post-hoc test, p < 0.05) in expression between the side and down configurations.

**Figure 8.**
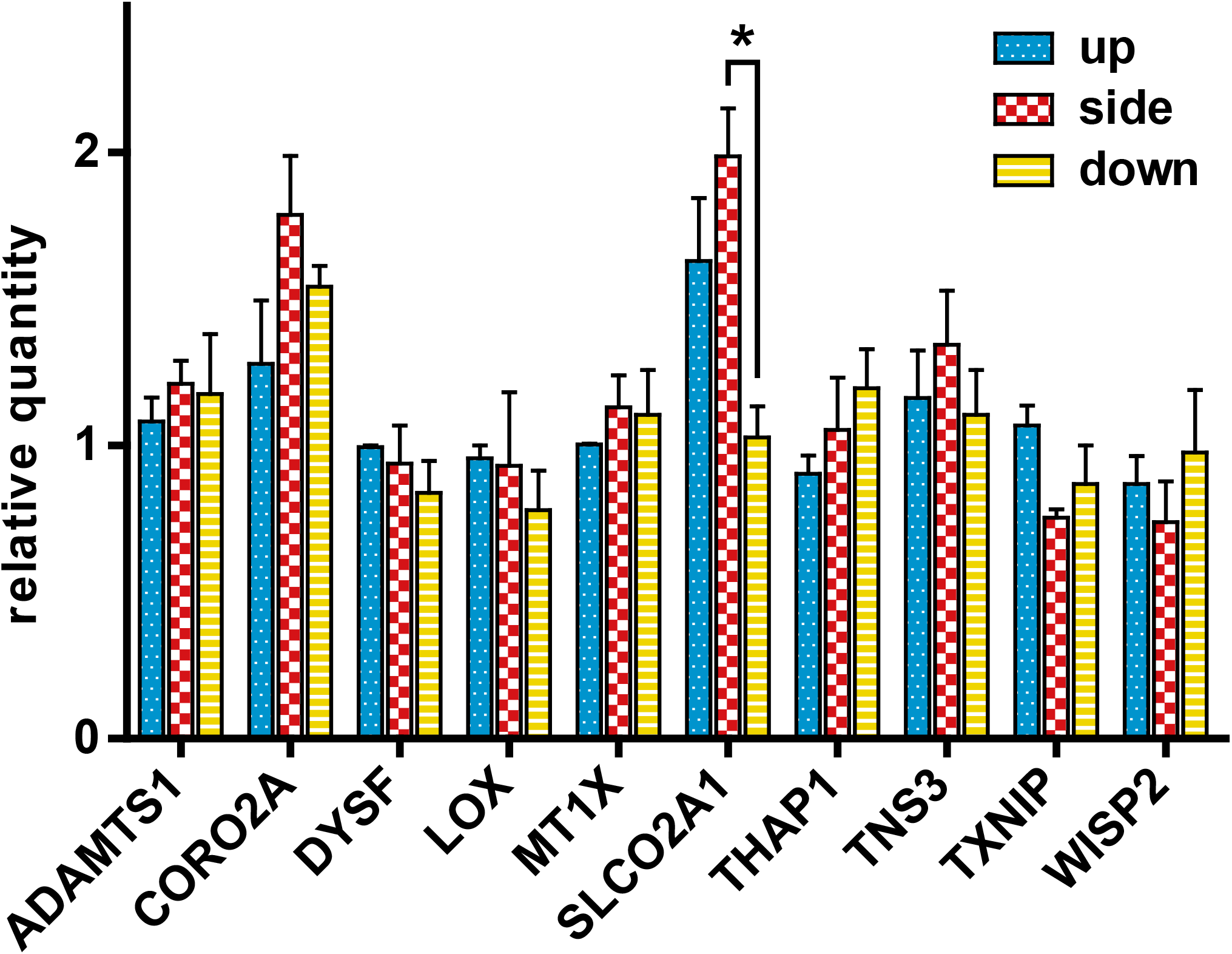
The c**omparison of** gene **expression** of the cells in the **up, side**, and **down configurations. The relative quantity of each gene was assayed using** qRT-PCR **and normalized to the 18S rRNA level. The data were expressed as the mean ± S.E.M. *: p < 0.05**.

## 4 Discussion

Researchers have used different types of species, organs, and conditions of gravity to evaluate microgravity’s effect. However, it is challenging to extract meaningful knowledge from these results. In this study, we performed a meta-analysis of gene expression using DNA microarray datasets obtained from experiments performed on diverse samples and species using various methods to mimic or achieve microgravity.

Interestingly, the t-SNE analysis revealed that gene expression patterns were grouped in DNA array datasets. For example, the datasets (G, H, K, and L) obtained from the skeletal muscle of space-flown mice formed a different cluster from datasets (A, B, C, D, E, and J) from skeletal muscle under simulated microgravity, except that one dataset from the space-flown sample (I) was in the “simulated microgravity” cluster and another dataset from simulated microgravity (F) was in the “spaceflight” cluster (Figs. 4B–D). Meanwhile, the datasets from hind limb unloading (D and E) were located far from the “spaceflight” cluster. This tendency was also observed when the number of analyzed genes was reduced to 177 (Figs. 4F–H). However, it should also be noted that the difference in experimental conditions between the datasets G/H/K/L and the datasets D/E was recognized not only in the mode of microgravity but also in the duration of microgravity and the method of sample storage. Nevertheless, according to the t-SNE analysis, the gene expression pattern in datasets D and E differed from G, K, and L. This fact was also supported by the result of cluster analysis (Fig. 4I).

In addition, the DNA array datasets obtained from endothelial cells under actual (X) and simulated (Y) microgravity are located close to each other (Figs. 4F-H), suggesting a remarkable similarity in gene expression patterns, given that the datasets come from independent research groups. This result provides a valid basis for estimating gene expression change under microgravity in space with simulated microgravity datasets in endothelial cells.

Next, we investigated the genes whose expression may be affected by microgravity. The study results demonstrated that *SLCO2A1* expression was decreased in both space-flown HUVECs (FC value = −0.23) and clinorotated EA.hy926 cells (FC value = −2.66) (Fig. 2**A**). Furthermore, the qRT-PCR analysis revealed that *SLCO2A1* expression was decreased in response to a 7-day exposure to SMG using a clinostat, in parallel to the results in space-flown HUVECs and clinorotated EA.hy926 cells (Fig. 6). Importantly, the decreased expression of *SLCO2A1* was not found in previous studies using space-flown HUVECs [7] and clinorotated EA.hy926 cells [17]. In other words, the decreased expression of *SLCO2A* has been identified via a meta-analysis of multiple sets of previously published data.

Other genes responsive to microgravity have also been uncovered, such as LOX, encoding LOX with a copper-dependent amine oxidase activity and catalytic activity in the cross-linking of collagen and elastin [20], was affected. *LOX* expression was increased, consistent with the results in EA.hy926 cells [17]. However, the expression of LOX is reduced in space-flown HUVECs [7]. Such discrepancy may be due to the differences in cell types and the conditions of microgravity, spaceflight vs. clinorotation. Another gene whose expression is altered by gravity changes is *LRP4*, which is involved in forming and maintaining neuromuscular junctions [21]. Interestingly, the expression of *LRP4* was increased in this meta-analysis (Fig. 3). It is tempting to speculate that one of the mechanisms of muscle atrophy under microgravity is the altered expression of LRP4; this hypothesis should be carefully studied in the future.

On the other hand, we analyzed the genes with identified sensitivity to gravity in the context of skeletal muscle but observed no changes in their expression in HUVECs exposed to SMG. When we analyzed the genes that showed sensitivity to gravity in skeletal muscle, no changes were observed in the gene expression level in HUVEC underwent SMG (Fig. 6B). This result suggests that modulation of gene expression is organ-specific. This is also suggested because the two DNA array datasets from endothelial cells are located outside of the muscle cluster in the t-SNE analysis.

Furthermore, the direction of gravity was found to affect the morphology, proliferation, and gene expression of HUVECs in this study. The decrease in the cell number in the “down” configuration is likely due to the change in the cell adhesion in response to gravity. However, the cells are unlikely to simply fall off, as the adhesion of extracellular matrix proteins to the culture flask is relatively stable. Cells are known to respond to gravity. In animals, the direction of gravity affects the proliferation of osteoblasts [22]. Interestingly, in our study, osteoblast proliferation is inhibited in the inverted culture, similar to the HUVECs cultured in the down configuration. On the other hand, plant cells respond to gravity’s direction, thus modulating the direction of root growth [23]. In plants, it is thought that the sedimentation of heavy statolith increases the tension of the actin cytoskeleton and subsequently activates mechanosensitive ion channels [24]. Suppose heavy nuclei and mitochondria also contribute to an animal cell’s gravity sensing mechanism similar to plant statolith. In that case, this mechanism may explain the change in gene expression depending on the direction of gravity observed in the current study.

Intriguingly, *SLCO2A1*’s expression was modulated in response to the direction of gravity in this study. In line with the results from previous studies using SMG, gravity appears to affect the expression of *SLCO2A1*, which encodes a prostaglandin transporter with a recently discovered involvement in the regulation of body temperature [25]. Although it is tempting to speculate that the altered thermoregulation in astronauts [26] is partly attributed to the altered expression of the *SLCO2A1*, this hypothesis should be tested in the future.

## Conclusion

Using the meta-analysis of DNA microarray datasets obtained from the vascular endothelial cells, we have identified gene candidates whose expression levels change under microgravity. Accordingly to the t-SNE analysis, among the mixed datasets from endothelial cells and skeletal muscle, the gene expression patterns in response to gravity in endothelial cells form specific clusters. Experiments using endothelial cells under simulated microgravity and different directions of gravity confirmed that the expression of one of the candidate genes, *SLCO2A1* encoding the prostaglandin transporter, is indeed altered. These results suggest that meta-analysis of multiple microarray data is useful for identifying novel target genes.

## 6 Author Contributions

KT and KN contributed to the conception and design of the study; YIL and MW performed the experiments; MW, YIL, YUL, CW, and KT wrote sections of the manuscript. All authors contributed to manuscript revision, and all authors read and approved the submitted version.

## 8 Abbreviations

FC: Fold change
HUVEC: Human umbilical vein endothelial cells
SMG: Simulated microgravity
t-SNE: t-distributed stochastic neighbor embedding

## 9 Acknowledgments

The authors gratefully acknowledge Central Research Laboratory, Okayama University Medical School for the assistance of qRT-PCR, Dr. Hitoshi Osada for preliminary research of this work, and Dr. Masatoshi Morimatsu for his valuable suggestions.

